# Estimation of task-based modulations in functional connectivity with MEG: a comparison of methods

**DOI:** 10.1101/407213

**Authors:** Juan L.P. Soto, Karim Jerbi

## Abstract

For the assessment of functional interactions between distinct brain regions there is a great variety of mathematical techniques, with well-known properties, relative merits and shortcomings; however, the methods that deal specifically with task-based fluctuations in interareal coupling are scarce, and their relative performance is unclear. In the present article, we compare two approaches used in the estimation of correlation changes between the envelope amplitudes of narrowband brain activity obtained from magnetoencephalography (MEG) recordings. One approach is an implementation of semipartial canonical correlation analysis (SP-CCA), which is formally equivalent to the psychophysiological interactions technique successfully applied to functional magnetic resonance data. The other approach, which has been used in recent electrophysiology studies, consists of simply computing linear correlation coefficients of signals from two experimental conditions and taking their differences. We compared the two approaches with simulations and with multi-subject MEG signals acquired during a visuomotor coordination study. The analyses with simulated activity showed that computing differences in correlation coefficients (DCC) provided better discrimination between true coupling changes and spurious effects; on the other hand, SP-CCA resulted in significant effects around the reference location which were not found with DCC, and which may be due to field spread. Based on our findings, we recommend the use of DCC for the detection of task-based changes in connectivity, as it provided better performance than SP-CCA.

## 1 Introduction

Electro- and magnetoencephalography (EEG and MEG, or E/MEG) are among the most popular medical imaging modalities in studies of brain function and behavior. Especially when compared with modalities such as functional magnetic resonance (fMRI) and positron emission tomography (PET), the ability of E/MEG to measure direct electrical brain activity, coupled with their high temporal resolution (on the order of milliseconds) makes them an attractive option for the functional neuroimaging community. The latter property, in particular, allows an accurate analysis in both the time and the frequency domains of the estimates of electric current densities.

One of the problems in Neuroscience to which E/MEG have been successfully applied, and that has been drawing increasing attention among researchers within the community, is functional connectivity, or the detection of interactions between spatially separated brain regions caused by a given experimental condition (Brookes et al., 2014b; Schoffelen and Gross, 2014; Hassan et al., 2014). Connectivity investigations based on EEG or MEG data usually assess interactions between two areas in the brain by comparing a single quantity from each area, such as amplitude envelope or phase within narrow, well-defined frequency bands, which is obtained from spectral or spectro-temporal representations of brain current density estimates. Some of the interaction measures implemented in recent findings dealing with functional connectivity are: coherence, or the normalized cross-spectral density at a given frequency (Gross et al., 2001; Nolte et al., 2004; Hipp et al., 2011); phase synchrony, or the average difference between the instantaneous phases of two narrowband signals (Lachaux et al., 1999; Palva et al., 2005; Fell and Axmacher, 2011; Belluscio et al., 2012); nested oscillations, or the coupling between the amplitude of a high-frequency time series and the phase of a low-frequency one (Canolty et al., 2006; Tort et al., 2010); and the correlation between the power or the amplitude of time series within specific frequency bands (Liu et al., 2010; Brookes et al., 2011). Regarding amplitude or power correlations, a recent methodological development involves the simultaneous comparison of brain activity at several frequencies between two locations using techniques from multivariate statistical inference (Soto et al., 2010; Brookes et al., 2012, 2014a), which not only take advantage of the increased sensitivity of multivariate analysis (Huberty and Morris, 1989), but also use some of its properties to discard spurious correlations caused by field spread (Schoffelen and Gross, 2009).

Although reports on brain states during rest are becoming more and more popular [e.g. de Pasquale et al. (2010); Boersma et al. (2011); Lansbergen et al. (2011); Hillebrand et al. (2012); Marzetti et al. (2013)], typical functional connectivity studies look for correlations between different areas in the brain during a specific cognitive task (visual, motor and so on), sometimes using as subjects patients suffering from a given condition, such as autism or epilepsy. In these studies, if one is interested in verifying the effects in connectivity that can be exclusively attributed to the task (or condition) under analysis, data can be acquired during rest (or from control subjects), and the comparison between the two experimental conditions can be performed either qualitatively (Srinivasan et al., 2007; Hall et al., 2013; Travis and Wallace, 1999) or with straightforward ANOVA models (Tal et al., 2013; Gootjes et al., 2006; Betti et al., 2013; Vardy et al., 2011; Bartolomei et al., 2006; Wada et al., 1998; Peters et al., 2013; Bettus et al., 2008; Jiang and Zheng, 2006; Peled et al., 2001; Cao and Slobounov, 2010). While these approaches for comparing conditions are capable of measuring task- or condition-based modulations in coupling, they do not account for other effects that might also be detected as fluctuations in correlations (such as variations in signal amplitude across conditions), which might result in spurious effects.

In Soto and Jerbi (2015), we presented a procedure to find condition- or task-based modulations in functional connectivity in time series obtained from MEG data, while minimizing the sources of spurious interactions mentioned in the previous paragraph. Our method was based on psychophysiological interactions, or PPI (Friston et al., 1997; Gitelman et al., 2003; O’Reilly et al., 2012), a technique that has been successfully applied in the computation of coupling changes from fMRI data. PPI implements a multivariate analysis-of-covariance model (MANCOVA), whereby the degree of linear relationship between two observation matrices **X** and **Y** is estimated, allowing a third matrix **Z** (Seber, 1984; Worsley et al., 2004); for the purposes of this study, observations **X** and **Y** refer to two different spatial locations in the brain, and the data for these observations come from current density power estimates at well-defined frequency bands, obtained from several repetitions of a MEG experiment consisting of two conditions (e.g. task and rest); these matrices are arranged in such a way that the coupling between them will be high if there is an increase or decrease of functional connectivity between the two locations from one condition to another. Observation **Z**, on the other hand, represents factors that might lead to a spurious coupling between **X** and **Y**, such as fluctuations in current density strength and signal leakage (O’Reilly et al., 2012). In preliminary tests with simulated and with real MEG recordings, PPI showed the potential of being a powerful tool for the analysis of task-based modulations in connectivity with E/MEG.

In this work, we extend our investigation of the performance of PPI in the detection of coupling changes with MEG signals, directly comparing it with univariate techniques that have been recently applied to the same problem – e.g. by (Brookes et al., 2016). We perform the comparison of our multivariate method with univariate techniques by means of simulated current densities and of real MEG data acquired during a visuomotor coordination study (Jerbi et al., 2007). The relative merits of both techniques will be ascertained based on their ability for finding regions with statistically significant coupling modulations and for identifying the signal frequency bands responsible for these strong effects.

## 2 Methods

### 2.1 Data collection

Let us consider that MEG data are acquired while a subject performs an experiment with two conditions: the task condition and the rest or control condition – generalizations for more than two conditions can be formulated based on analogous methods dealing with PPI (McLaren et al., 2012). The experiment is repeated several times for each condition: the number of repetitions (trials) for condition 1 is *n*_trials,1_, and for condition 2 is *n*_trials,2_. The *i*^th^ trial from the *j*^th^ condition – i.e. the *i*^th^ repetition of the experiment for condition *j, j* ∈ {1, 2} – will result in a data array **M**_*ij*_ (*n*_sensors_ × *n*_timepoints_) whose rows contain the time series of the measured magnetic fields at each MEG sensor. The relationship between **M**_*ij*_ and the brain activity array **S**_*ij*_ (*n*_sources_ × *n*_timepoints_), assumed to consist of elemental electric current sources (dipoles) constrained to the cortical surface and normally oriented to it, is given by the following linear equation:

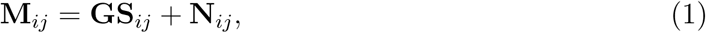

where **G** (*n*_sensors_ × *n*_sources_) is known as the forward operator or lead field matrix, and **N**_*ij*_ represents noise. The lead field maps the brain current densities onto the sensor space, and depends on the conductivity and geometry properties of the head; it can be estimated with simplified spherical head models or with more realistic finite element methods (Wolters et al., 2006; Mosher et al., 1999). There are several methods to obtain an estimate of **S**_*ij*_ from matrices **M**_*ij*_ and **G**; here, we implement a Tikhonov regularized minimum-norm inverse solution (Tikhonov and Arsenin, 1977; Okada, 2003):

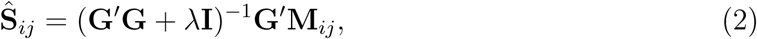

where *λ* = 10^−7^ is a regularization parameter, **I** is the identity matrix, and **G**′ denotes the transpose of matrix **G**. Matrix **Ŝ**_*ij*_ is made of row vectors 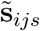 representing the estimated current density time series at each location *s* in the brain. We chose regularized minimum-norm imaging to compute **Ŝ** because its implementation is straightforward and because it requires no assumptions with respect to the number of active brain locations, their size, and how they interact; however, the methods described here are appropriate to any inverse solution.

In this study, we are interested in measuring changes in interaction of signal energies within specific frequency bands between distinct brain regions. With this goal in mind, we compute the power spectral density (PSD) of the current density estimate 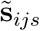 at each spatial location, then average the PSD over six frequency bands: delta (2 – 4 Hz), theta (5 – 8 Hz), alpha (8 – 12 Hz), beta (15 – 30 Hz), low-gamma (30 – 60 Hz) and high-gamma (60 – 90 Hz). We chose these bands due to the functional role they play in the somatosensory cortex (Jerbi et al., 2004), which is expected to be active in tasks such as the one carried out by subjects during the acquisition of the real MEG data used here (more details below). Thus each time series 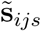 results in a row vector **p**_*ijs*_ with length *n*_freqs_ = 6 containing the energy content of 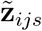 at the six frequency bands of interest. These vectors **p**_*ijs*_ will be the inputs to our SP-CCA model.

### 2.2 Semipartial canonical correlation analysis

For ease of notation, we will omit the index *s* from now on (e.g. **p**_*ijs*_ = **p**_*ij*_), since the SP-CCA model will be applied to all spatial locations in the brain. Further, one of the locations in the brain is chosen as the reference location, and the activity at this reference will be compared to that at all other locations. For a given location, we concatenate all energy estimates from condition 1 (i.e. all **p**_*i*1_) to form matrix **P**_1_, and matrix **P**_2_ is formed by concatenating all **p**_*i*2_. In a similar fashion, we create matrices **Q**_1_ and **Q**_2_ containing of the energy estimates of the reference location for each condition. The observation matrices used in SP-CCA are thus:

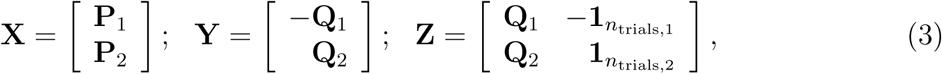

where **1**_*N*_ is defined as a *N* × 1 matrix whose elements are all ones. In our method, we examine the linear relationship between **X** and **Y**, controlling for effects on **Y** of a third matrix **Z**. Matrices **X** and **Y** are arranged in such a way that, for instance, an interaction measure computed from them will be high if a given frequency pair (one from each location) is strongly correlated in one condition, but strongly anti-correlated in the other. On the other hand, the configuration of matrix **Z** seeks to attenuate effects that also yield high interaction measures but that do not result from actual modulations in coupling: the term 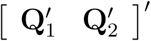 deals with strong correlations between the two locations that do not vary from one condition to another, while the term 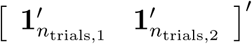 compensates strong fluctuations in signal power (O’Reilly et al., 2012).

The maximum semi partial canonical correlation between **X** and **Y**, controlling for **Z**, is given by the expression:

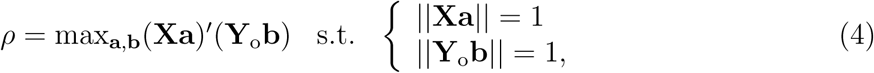

where matrix **Y**_o_ is the projection of **Y** onto the orthogonal complement of the subspace spanned by **Z**. The solution to this maximization problem yields the following eigenvalue equations:

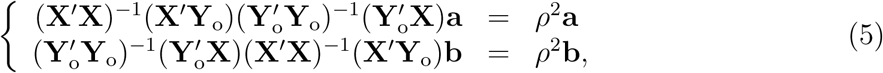

thus the maximum eigenvalue of either 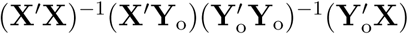 or 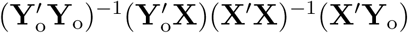 is the maximum semipartial canonical correlation *ρ*, and the corresponding eigenvectors, with size *n*_freqs_ × 1, are known as the canonical vectors **a** and **b**.

### 2.3 Difference of correlation coefficients

Let 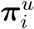 and 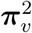 be, respectively, the *u*^th^ and *v*^th^ columns of matrices **P**_1_ and **P**_2_ described in the previous section; similarly, 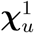 and 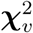 are the *u*^th^ and *v*^th^ columns of matrices **Q**_1_ and **Q**_2_ (*u, v* ∈ {1, 2,*…, n*_freqs_}). Then, in order to estimate the degree of task-based changes in coupling between two locations, we can compute the following statistic:

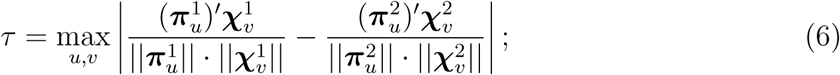

in other words, we calculate, for all possible combinations of frequency bands in the two locations under study, the difference between their (univariate) correlation coefficients for both conditions, and take the maximum absolute value (Brookes et al., 2016). For the remainder of this article, this method to assess interaction modulations will be called Difference of Correlation Coefficients, or DCC.

### 2.4 Statistical thresholding

Applying the procedures described in subsections 2.2 and 2.3 to all locations in the brain (the SP-CCA and DCC methods, respectively), we will have a brain map of *ρ* (or *τ*) statistics, reflecting the degree of coupling changes between the reference and every other spatial location. For either procedure, the first step to determine the locations with a statistically significant effect is to convert the *ρ* or *τ* map into a map of p-values; each p-value is obtained from the probability distribution of the computed statistic at the desired location, and we find this distribution empirically with a nonparametric resampling method (Nichols and Holmes, 2002; Pantazis et al., 2005). For the distribution of *ρ*, we create surrogate observation matrices that are statistically equivalent to the original **X**, **Y** and **Z** shown in equation (3) under the null hypothesis 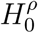 of no coupling modulations (which also implies that *ρ* = 0 under 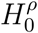), obtain from the new matrices a surrogate statistic 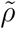, and repeat these steps several times; the histogram of all surrogate 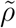 plus the original *ρ* will form the empirical probability distribution for the statistic. In our case, the surrogate models are constructed by randomly shuffling the rows of matrix **X**, keeping **Y** and **Z** unchanged. We proceed in a similar fashion to obtain the empirical distribution of *τ,* using equation (6) to compute the surrogate statistics 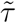 but with matrices 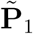 and 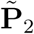 instead of **P**_1_ and **P**_2_ (observations **Q**_1_ and **Q**_2_ are not altered). We create 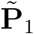 by randomly shuffling the rows of matrix 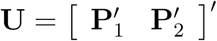 and taking the first *n*_trials,1_ rows – 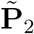 is then given by the remaining *n*_trials,2_ rows of the randomized **U**.

A location will have significant changes in interaction with the reference if its p-value *p* is lower than a threshold *p*_*α*_. The choice of *p*_*α*_ must take into account the very large number of statistical tests to be performed (as many as the number of locations in the brain), which can inflate the number of false positives. The procedure we implement to compensate for this undesired effect seeks to control the familywise error rate (FWER), or the probability of at least one false positive among all locations under *H*_0_ (Nichols and Hayasaka, 2003). The FWER thresholding procedure for a confidence level (1 – *α*) (throughout this work, *α* = 5%) consists of the following steps (Holm, 1979; Hochberg, 1988):

1. Order the location p-values from smallest to largest:

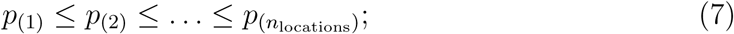
2. Starting at *k* = 1, verify the following inequality, for increasing values of *k*:

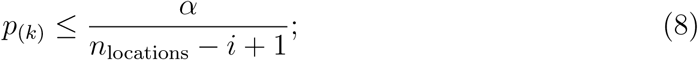
3. Once a p-value *p*_(*K*)_ is found for which the inequality above is violated, declare all locations corresponding to the p-values *p*_(1)_, *p*_(2)_,*…, p*_(*K*–1)_ statistically significant – i.e. *p*_*α*_ = *p*_(*K*)_.

### 2.5 Contribution of individual frequencies to detected effects

Let us suppose that, having determined that there is a statistically significant change in coupling between two spatial locations according to the SP-CCA method, we are interested in verifying whether frequency band *i* at the reference location and frequency band *j* at the other location contribute strongly to the detected effect. If **x**_*i*_ and **y**_*j*_ are, respectively, the *i*^th^ and *j*^th^ columns in matrices **X** and **Y**_o_ (used to compute the *ρ* statistic, according to equation 5), then we can perform this verification simply by computing the absolute value of the correlation coefficient between **x**_*i*_ and **y**_*j*_:

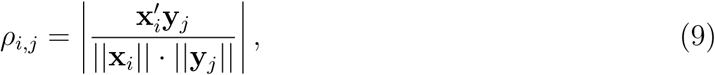

which is then compared with *ρ*_*α*_, the value of *ρ* corresponding to *p*_*α*_ (i.e. the p-value used for the FWER-correcting procedure described in subsection 2.4). (Carbonell et al., 2009; Soto et al., 2016). If the DCC method was implemented instead of SP-CCA, the approach is similar: for a given pair of frequency bands {*i, j*} at each location, we first find *τ*_*i,j*_:

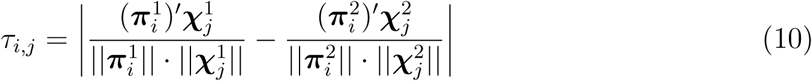

(i.e. the argument of the maximum operator in equation 6), and afterwards check whether *τ*_*i,j*_ ≥ *τ*_*α*_ (the value of *τ* corresponding to *p*_*α*_).

## 3 Results

We employ synthetic MEG signals to compare the performances of SP-CCA with that of DCC in terms of detection accuracy and robustness to noise, and illustrate the practical implications of the choice of either method with MEG data obtained from 15 subjects during a visuomotor coordination study – in Jerbi et al. (2004, 2007); Besserve et al. (2007) this experiment is discussed in more detail. Briefly, the two conditions tested were: task (VM), where the subjects attempted to control with a trackball the random rotations of a cube appearing on a screen in front of them; and rest (R), where the subjects only looked at a screen where a motionless cube appeared, and did not operate the trackball. The data were recorded during eight-minute runs, where the conditions were switched every 8-to-12 seconds; the signals acquired during these runs that were not discarded due to movement artifacts were then corrected for heart beat effects and divided into 1-second trials, which were the inputs to our SP-CCA and DCC analyses (the average number of trials across subjects was 216 for the VM condition and 107 for R). The inverse procedures and time-frequency analysis used here were performed with the open source software package BrainStorm (**?**).

### 3.1 Proof of concept: simulations with Gaussian noise

The simulated brain activity **S**_*ij*_ used to compare SP-CCA with DCC had the spatial profile shown in figure 1a. It consisted of four sources, two in the left hemisphere and two in the right. Time courses in each source obeyed the following equations:

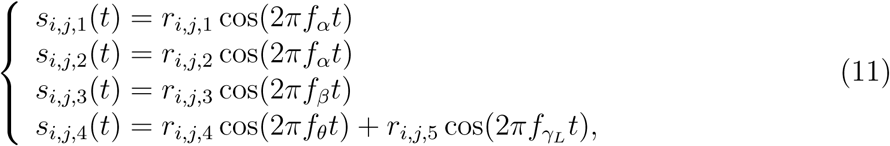

where *f*_*θ*_ = 6.5Hz, *f*_*α*_ = 10Hz, *f*_*β*_ = 22.5Hz, *f*_*γL*_ = 45Hz and 0 ≤ *t* ≤ 1s. The squares of amplitudes of the cosine waves *r*_*i,j,n*_, *n* ∈ 1,*…,* 5, vary randomly across trials and are sampled from a multivariate Gaussian distribution with mean *µ*′ = [2 2 2 2 2]′ and covariance matrices Σ_1_ and Σ_2_ depending on condition:

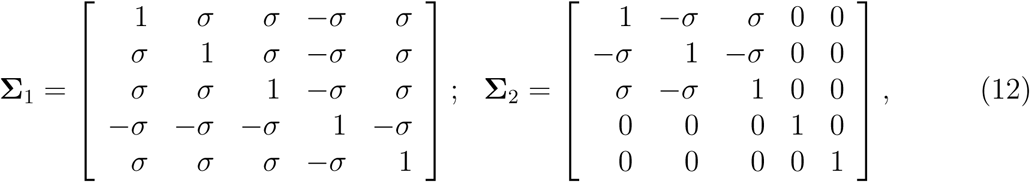

for *σ* = 0.9. Among other effects, this simulation setting implies that there is alpha-alpha positive correlation between sources 1 and 2 in one condition, which becomes negative correlation in the other condition; that alpha and beta are correlated between sources 1 and 3, and the strength of the coupling does not change across conditions; and that there is negative alpha-theta and positive alpha-low-gamma coupling between sources 1 and 4 only in one of the two conditions.

**Figure 1:**
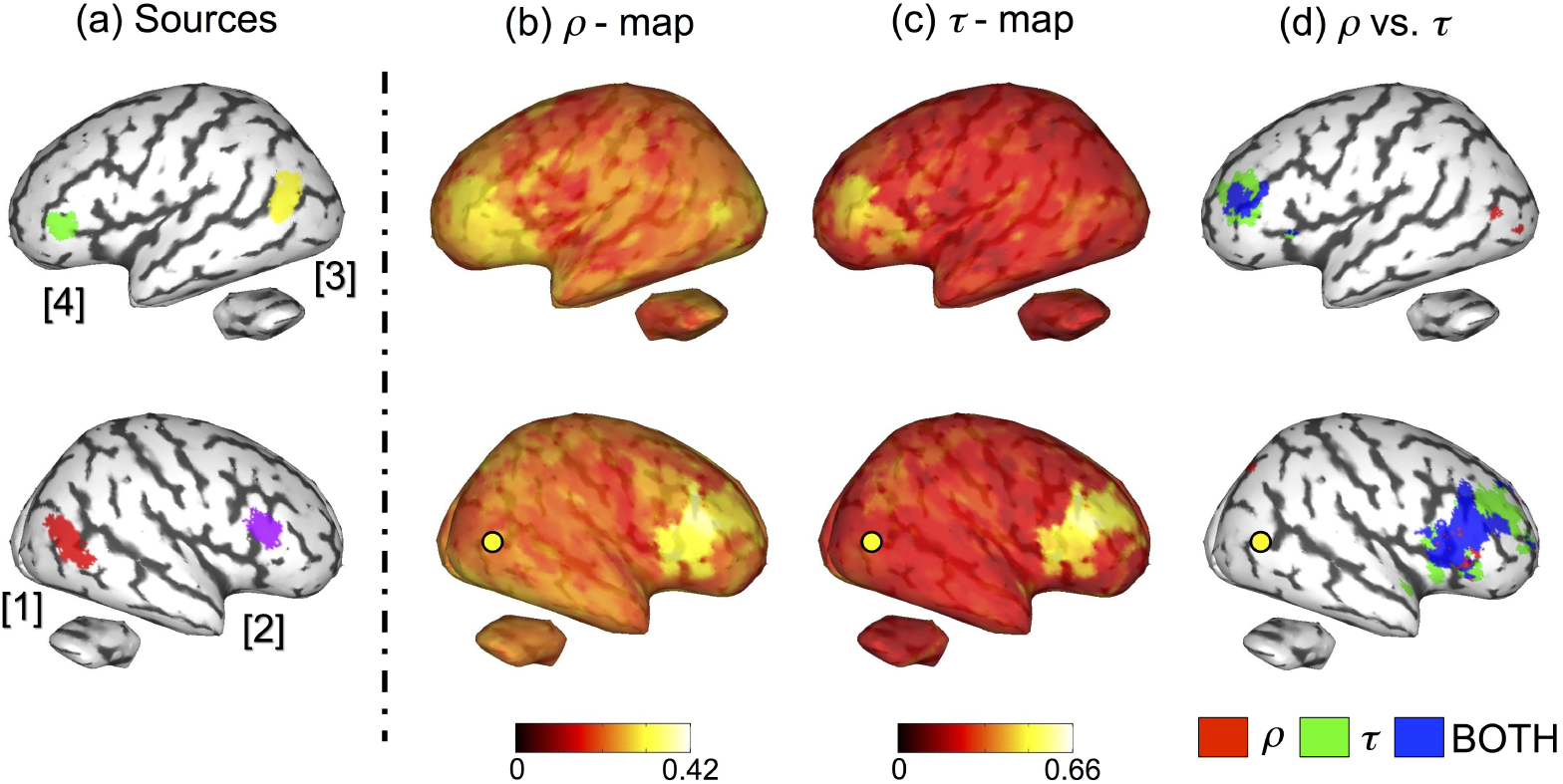
Simulation results. (a) The spatial profile of the four simulated sources; the signals in regions 1 and 2 lied in the alpha band, the signal in region 3 was within the beta band, and there was theta and low-gamma activity in region 4; (b) Map of the SP-CCA statistic *ρ* when the reference is in region 1 (indicated by the yellow dot); (c) Map of the DCC statistic *τ* when the reference is in the region 1; (d) Map of statistically significant locations obtained with SP-CCA only (red locations), DCC only (green locations), and both methods (blue locations), with reference in region 1.

We created 200 activity matrices **S**_*ij*_ for each condition, which were then projected into the sensor space according to equation (1), and contaminated with Gaussian noise, with mean zero and standard deviation 2 × 10^−3^ (which represents noise with energy 100 times higher than the mean energy of the simulated sources, when projected onto the sensor space). Finally, we obtained estimates of the current density maps with the inverse operator (equation (2)), created observation matrices as described in section 2.1, and computed *ρ* and *τ* maps using the SP-CCA and DCC procedures, respectively (subsections 2.2 and 2.3). The results of our computations are presented in figures 1b-1d: in figure 1b, we can see the values of *ρ* for every spatial location in the brain when the reference location lies within source 1; similarly, figure 1c presents a brain map of values of *τ* with the same reference as in figure 1b; and figure 1d displays the brain locations with statistically significant values of *ρ* and of *τ,* based on the thresholding procedure described in section 2.4 According to these maps, both methods were able to identify task-based variations in coupling between the reference and regions around sources 2 and 4; these findings were consistent with our simulation settings, since alpha-alpha coupling between sources 1 and 2 changed from positive (in condition I) to negative (in condition II), and there was alpha-theta and alpha-low-gamma coupling between sources 1 and 4 only in condition (I). However, while DCC yielded no other significant activity, implementation of SP-CCA resulted in a few active locations near source 3, where there was alpha-beta coupling with the reference that did not vary with condition.

To find out the frequency bands at the reference and at the other locations that caused the detected effects shown in figure 1d, we applied the post-hoc analysis discussed in section 2.5 for the SP-CCA and the DCC approaches; we display the results of these tests in figure 2. According to these maps, DCC was able to identify correctly the frequency bands involved with the detected interaction modulations, and their signals (i.e. alpha-theta decrease and alpha-low-gamma increase between sources 1 and 4, and alpha-alpha decrease between sources 1 and 2), but it also incorrectly found an alpha-theta contribution to the significant effect detected in region 2, possibly due to signal leakage. On the other hand, SP-CCA only identified an alpha-alpha decrease in coupling between sources 1 and 2.

**Figure 2:**
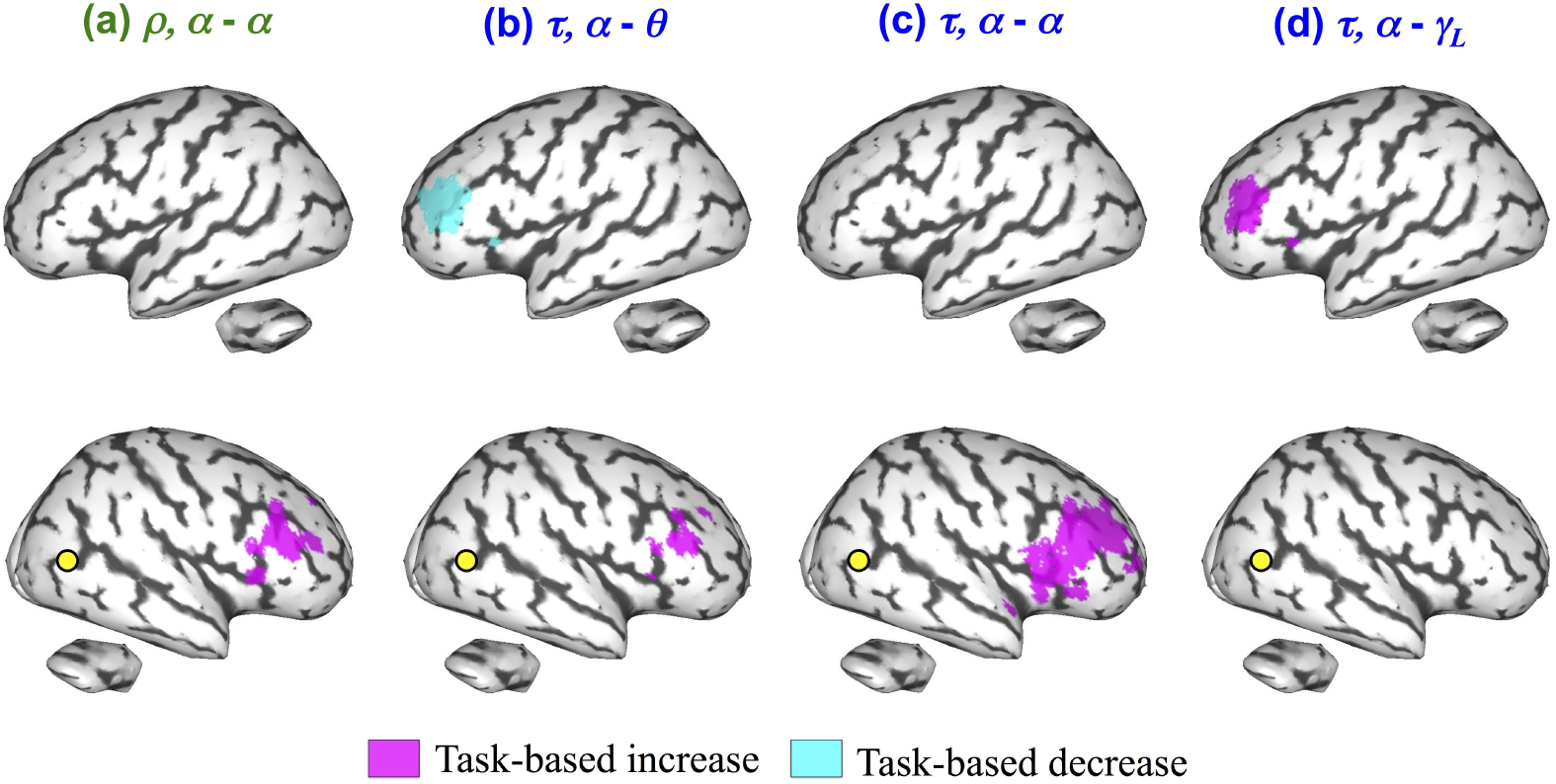
Brain maps, obtained from the simulated data used for the images in figure 1, of spatial locations where there was a significant change in alpha-alpha coupling with SP-CCA [column (a)], and in alpha-theta, alpha-alpha, and alpha-low-gamma with DCC [columns (b), (c) and (d), respectively] (no other frequency band pairs yielded significant contributions to the detected effects with either method). Purple locations indicate an increase in coupling, and blue regions represent a decrease in coupling.

In short, figures 1 and 2 indicate that DCC appears to be more reliable than SP-CCA with respect to robustness against false positives, as only the latter identified significant regions around source 3 (where there was no change in correlation), even though neither method could single out accurately the frequency bands that caused significant changes in coupling (DCC detected alpha-theta contributions around source 2, and SP-CCA did not find any frequency responsible for the effects near source 4). In the next subsection, we investigate more thoroughly the comparative performances of SP-CCA and DCC by means of ROC analyses.

### 3.2 ROC analysis: simulations combined with experimental data

We further investigated the performance of SP-CCA with respect to detection accuracy and robustness to noise with an ROC analysis – specifically, a method adapted from a variant of ROC called free-response ROC (FROC) (Egan et al., 1961; Metz, 2006). As in subsection 3.1, we use simulated brain sources, but instead of Gaussian noise, we attempted to emulate the noise properties usually found in MEG measurements by adding to our synthetic data real human MEG signals, which act as background noise.

We started each of the ROC analyses performed here by creating 100 datasets, similar in structure to the dataset used for the proof of concept in section 3.1. The 100 datasets, each consisting of 200 MEG trials, were divided into two groups:

- *Group 1 datasets* had 100 trials belonging to condition (I) and 100 trials belonging to condition (II). In condition (I), there are two activity sources in the brain whose location was chosen randomly (with the condition that they be at least 5 cm apart from each other). The time series of the activity at the sources were cosine waves whose power varied across trials and followed a Gaussian distribution with mean 2, variance 1 and correlation coefficient between them *σ*; the frequencies of the cosine waves were chosen randomly among the central frequencies of the six bands of our study (as long as the two frequencies were different). The trials of condition (II) were created as the ones of condition (I), except that the frequency of the signal at the second source changed from the one it had in condition (I), while still being different from the frequency of the first source.
- *Group 2 datasets* also had 100 trials for condition (I) and 100 trials for condition (II), and they were created as the trials for the group 1 datasets, but in this case the signal frequencies at both sources remained the same across conditions – i.e. there was no difference in brain activity between the two conditions.

The real MEG measurements we added to the simulated sources (after projecting them onto the sensor level) came from a set of single-subject 231 trials acquired during the VM condition of the visuomotor experiment briefly described at the beginning of section 3; to each one of the 200 trials of simulated data (from any dataset), one randomly chosen MEG trial was added. These new trials containing both synthetic and real MEG signals were converted, first into the inputs to our SP-CCA and DCC models, then into *ρ* and *τ* maps (having as reference a location within one of the sources), as we did in section 3.1. Finally, we defined the true positive fraction (TPF) and the false positive fraction (FPF) for each statistic map, based on these 100 p-value maps and for a given value of *ϑ* (0 ≤ *ϑ* ≤ 1), according to the following:

- The TPF was given by the ratio of *group 1 datasets* where there was at least one spatial location within 2 cm of the center of the other source (i.e. the source not chosen as the reference) with *ρ* (or *τ*) greater or equal than *ϑ*;
- The FPF was given by the ratio of *group 2 datasets* where there was at least one spatial location anywhere in the brain with *ρ* (or *τ*) greater or equal than *ϑ*.

Computing the TPF and the FPF for several values of *ϑ* gave us the ROC curve for the corresponding datasets.

Figure 3 shows values of the areas under ROC curves (AUCs) obtained with the SP-CCA and the DCC methods, where the value of the correlation coefficient *σ* between the signals at the simulated sources was equal to 0.7 and for varying levels of the SNR – in this section, the square of the SNR was the ratio of the average energy of the simulated sources projected onto the sensor level (across trials, time and space) to the average energy of the 200 real MEG trials. Similarly, figure 4 displays AUC values for varying values of *σ* and for SNR equal to 1/100. According to these plots, DCC outperformed SP-CCA in almost all scenarios – the only exception was the case where SNR = 0.001, a very noisy situation in which both methods performed poorly. These results thus provide further evidence of the advantages of using an univariate approach to detect coupling modulations, in terms of its ability to discriminate between true brain activity and spurious effects.

**Figure 3:**
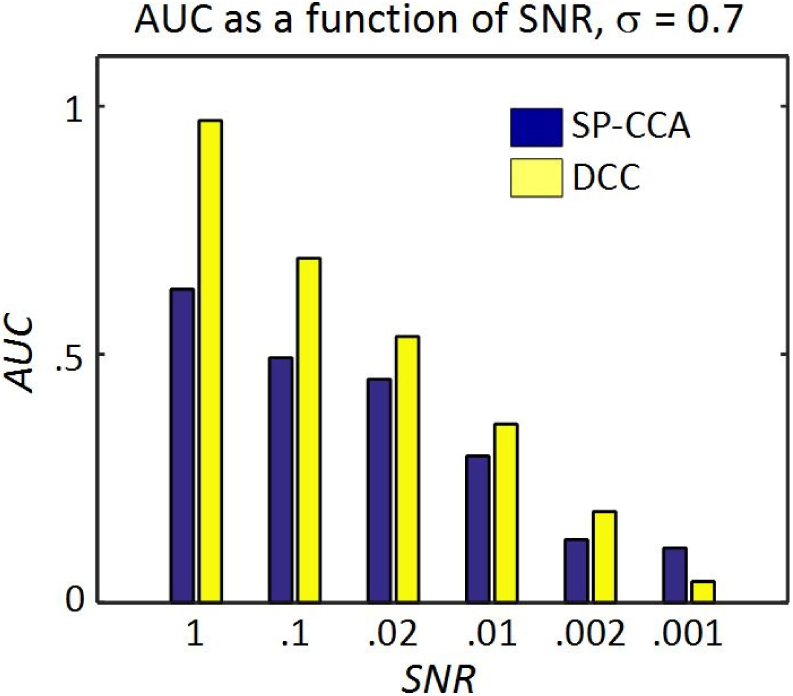
Values of the areas under ROC curves (AUCs) for different values of the SNR, and for *σ* = 0.7. Blue bars indicate SP-CCA, and yellow bars represent DCC.

**Figure 4:**
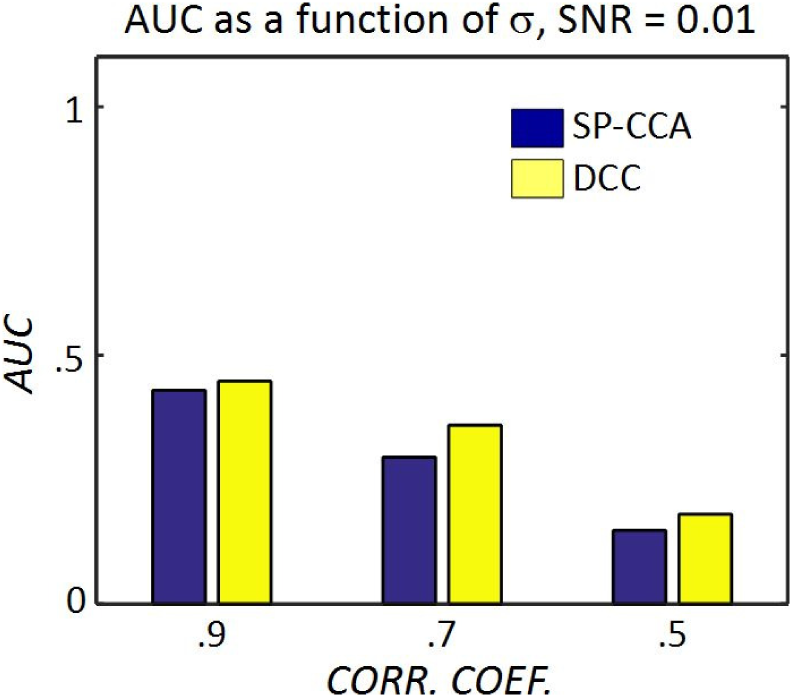
Values of the areas under ROC curves (AUCs) for different values of the correlation coefficient *σ*, and for SNR = 0.01. Blue bars indicate SP-CCA, and yellow bars represent DCC.

### 3.3 Experimental data

The results of the comparisons between SP-CCA and DCC with respect to the detection of coupling changes in the VM experiment when a reference was located in the left motor cortex are displayed in figure 5. In these images, the color coding for each spatial location indicates the number of subjects where the value of *ρ* (or *τ*) was statistically significant for that location: for instance, the highest value for a location in the maps on the left column was 12, which means that at most 12 out of the 15 subjects analyzed had a significant *ρ* (for the maps on the right column, on the other hand, the highest value across all locations was 5). Based on these maps, we may notice that, with either method, almost all the brain regions have a significant effect in at least one subject tested. Furthermore, although the discrepancies between the methods are not very strong for locations in the right hemisphere (i.e. the *ρ* and *τ* maps are active in about the same brain regions, and in comparable numbers of subjects), there is a marked difference between SP-CCA and DCC in the left hemisphere, particularly in the neighborhood of the reference location, where the number of subjects with an effect becomes noticeably larger with SP-CCA than that obtained with DCC.

**Figure 5:**
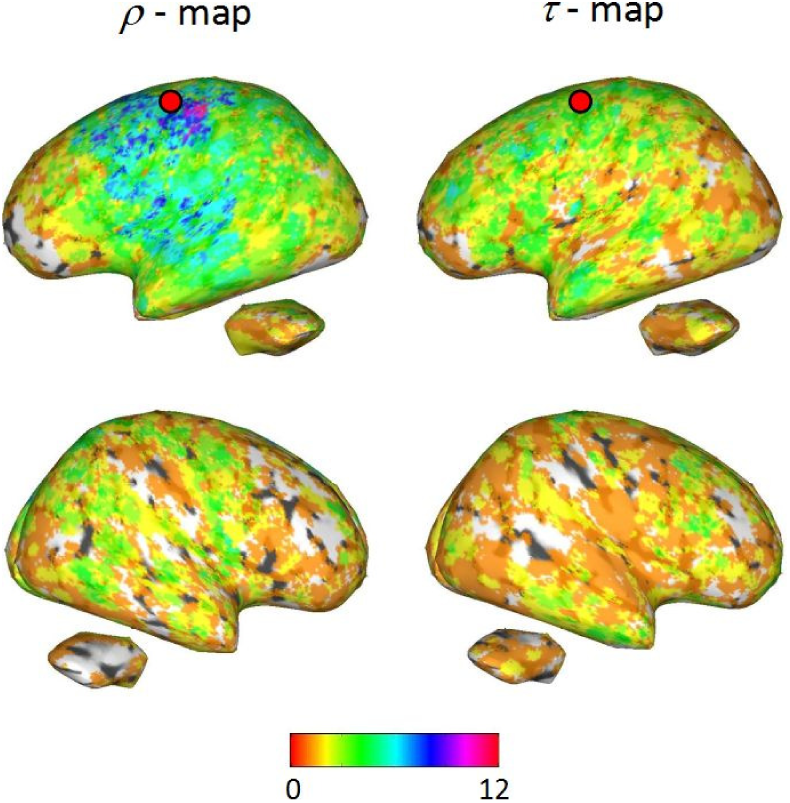
Multi-subject task-based modulations in coupling with the SP-CCA (left column) and the DCC (right column) methods, obtained from the VM data. For a given spatial location, its color represents the number of subjects where this location was statistically significant when *ρ* or *τ* was computed.

As for the frequency bands that are responsible for the identified effects, figures 6 and 7 show, for a selection of frequency band pairs, maps indicating how many subjects had significant contributions from the pair under consideration. Specifically, the pairs chosen for figure 6 are those that were active in more subjects than any other pair with SP-CCA and that are associated with a task-based *increase* in coupling, while the pairs that appear in figure 7 are also those present in the most subjects when SP-CCA is computed, but associated with a task-based *decrease*. In both cases, we can see that DCC tends to be more conservative than SP-CCA, as the active areas with the latter method are on the whole more widespread (the main exception being the *β – β* decrease found in the right hemisphere).

**Figure 6:**
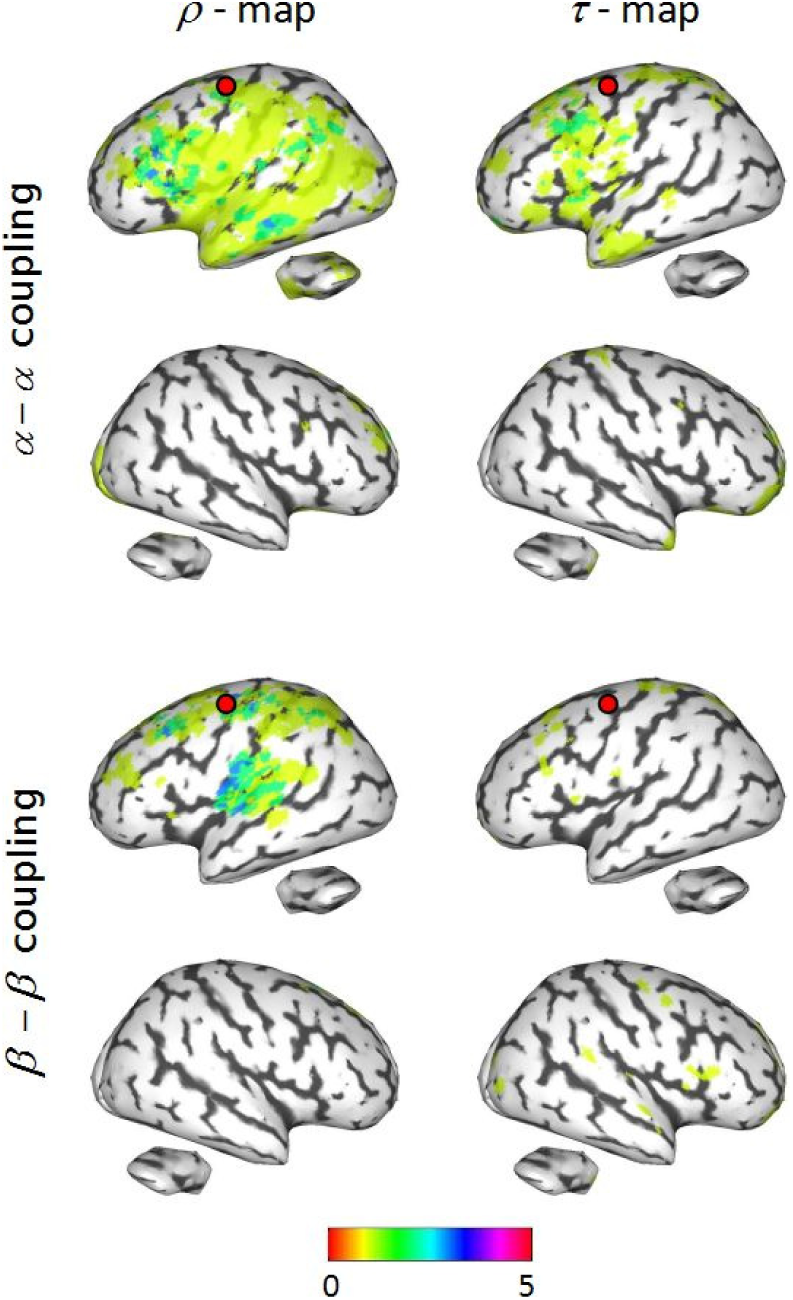
Brain locations where there the *α* – *α* (first and second rows) and *β* – *β* (third and fourth rows) frequency band pairs contributed significantly to task-based *increase* in coupling in at least one of the subjects analyzed during the VM experiment. The color coding of the spatial locations represents the number of subjects where there was a strong contribution of the frequency band pair under study.

**Figure 7:**
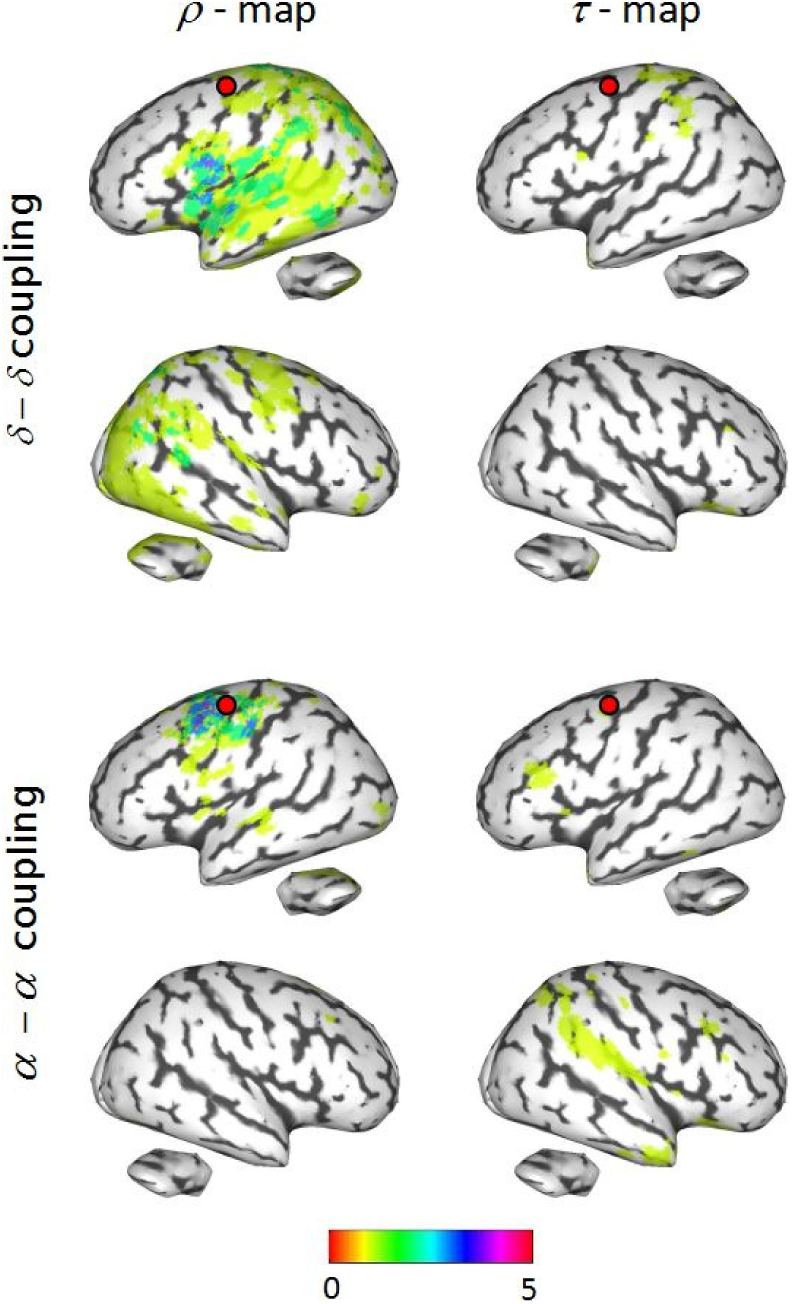
Brain locations where there the *δ* – *δ* (first and second rows) and *α* – *α* (third and fourth rows) frequency band pairs contributed significantly to task-based *decrease* in coupling in at least one of the subjects analyzed during the VM experiment. The color coding of the spatial locations represents the number of subjects where there was a strong contribution of the frequency band pair under study.

## 4 Discussion

In this work, we performed a comparison between two approaches to identify task-based modulations in functional interactions between two brain regions: semipartial canonical correlation analysis (SP-CCA), and difference of correlation coefficients (DCC). Our analyses with synthetic MEG signals showed that the latter method was more accurate in discriminating between true brain activity and spurious effects; as a consequence of this discrepancy in performance between the two methods, the use of SP-CCA may result in an excessive number of incorrectly detected brain locations, as our tests with data acquired during a visuomotor coordination study suggest. However, in order to ascertain that these locations detected by SP-CCA but not by DCC are in fact false positives, a more thorough investigation of the implications on brain activity of visuomotor coordination is necessary, possibly by means of tools with better spatial resolution (such as ECoG, depth electrodes etc.)

As our simulations showed, particularly the results in section 3.1, one of the disadvantages in using SP-CCA is its inability to identify brain locations where there are strong task-based coupling fluctuations. In a tutorial article dealing with PPI (which is simply SP-CCA with a different mathematical formulation), O’Reilly et al. (2012) explain that this lack of power is an unavoidable characteristic of the technique, because matrices **Y** and **Z** in equation 3 are correlated by construction, and because SP-CCA seeks to identify only linear relationships between **X** and **Y** that are not explained by **Z**; it is a drawback that does not affect DCC, since this approach simply takes into account the linear relationships between the corresponding columns of **X** and **Y**. O’Reilly et al. (2012) mention other possible sources of degradation in the performance of PPI, such as nonlinearities and subject behavior (response to errors, learning etc.); however, these are factors that may also affect the performance of DCC, and the extent to which they impact either method is beyond the scope of the present work.

A phenomenon that was not dealt with directly in our studies was spurious correlations due to linear mixing of sources, caused by the poor spatial resolution of brain activity reconstructed with E/MEG (Schoffelen and Gross, 2009). Here, we implicitly adopted the simplifying assumption that linear mixing was constant across conditions, thus with any method we employed to estimate differences in coupling, these artifacts would be canceled out. Although our simulations with Gaussian noise and simple, narrowband sources (subsection 3.1) indicate that both methods are able to deal adequately with the spurious coupling, the performance of the SP-CCA method in the ROC analysis (subsection 3.2) and the higher number of subjects with strong changes in coupling near the reference location yielded by SP-CCA (subsection 3.3) provide some evidence that this method may in fact not be as reliable as DCC in discarding interactions due to field spread – and in situations, one must remember, with brain activity and noise with much more complicated shapes and across wider frequency bands. In any case, one should be careful when including procedures to correct for linear mixing in the *ρ* and *τ* statistic maps, either due to methodological concerns (e.g. how the correction procedure is affected by different trial numbers between conditions) or because corrected brain maps lose in statistical power, since getting rid of linear mixing effects tends to also eliminate true brain activation (Nolte et al., 2004; Soto et al., 2016).

In our investigations with real MEG signals, we analyzed data from multiple subjects, but we did not execute any procedure to pool the data from them into a single group statistic map, with either method; instead, we obtained thresholded *ρ*- or *τ*-maps individually for each subject, and then simply counted, for any brain location, for how many of the 15 subjects that location was statistically significant. We proceeded thus because our main goal with the VM data investigations was not to establish a definite connection between brain electrophysiology and visuomotor coordination, but to demonstrate the differences between SP-CCA and DCC when actual MEG time series were manipulated. Furthermore, the statistical methods typically used for group analysis are not without interpretation issues: for instance, popular approaches such as the minimum p-value (Tippett, 1931) and the transformation formulas proposed by Fisher (1954) and by Stouffer et al. (1949) tend to be too lenient (since a single subject with a very low p-value is enough to result in a strong group effect), while more recent developments, where there is a strong multi-subject effect if it appears in a pre-determined fraction of the subjects (Wilkinson, 1951; Heller et al., 2007), are still dependent on a parameter whose choice of value is subjective. Even approaches such as Edgington’s method (Edgington, 1972), which in a sense deals with average effects, would not be of use for our data, especially in the tests of individual contributions of frequency bands, where no frequency band pair was significant in more than five of the fifteen subjects.

